# Is the predation risk of mate-searching different between the sexes?

**DOI:** 10.1101/363507

**Authors:** Viraj R. Torsekar, Kavita Isvaran, Rohini Balakrishnan

**Affiliations:** Centre for Ecological Sciences, Indian Institute of Science, Bangalore, India

**Keywords:** Communication, sex-specific costs, sex roles, predation risk, mate searching, crickets

## Abstract

In animals that communicate for pair formation, generally one sex invests more effort in mate searching. Differential predation risk of mate searching between the sexes is hypothesised to determine which sex invests more effort in mate searching. Although searching by males is prevalent in most animals, in orthopteran insects and some other taxa females physically move to localise signalling males who are predominantly sedentary. Although the two sexes thus share mate searching effort in orthopterans, their behavioural strategies are different and sexual selection theory predicts that signalling males may be following the riskier strategy and incurring higher costs. However, relative levels of risk posed by the two mate searching strategies remain largely unexplored. Hence, we estimated the relative predation risk experienced in natural populations by signalling males and responding females. We did this by quantifying predation risk as a probability of mortality in the context of acoustic communication in a tree cricket, *Oecanthus henryi* from its ecologically relevant predator, a lynx spider, *Peucetia viridans*. Spiders may perceive calling in males and movement in females by their ability to detect both airborne acoustic cues and substrate-borne vibratory cues. Probability of mortality was quantified by partitioning it into three spatial components at which crickets and spiders interact, using a combination of extensive field observations and manipulative experiments in a semi-natural setup. We found no differences in predation risk faced by calling males and responding females, supporting the prediction that similar sex-specific costs can explain shared mate searching responsibilities. Our findings therefore suggest that direct benefits offered by males to females upon pair formation may better explain shared mate searching effort between the sexes in orthopterans.

## Introduction

Searching for mates typically involves some physical activity for pair formation. Mate searching effort is defined “as a costly investment in traits that facilitate encounters with potential mates, including mobility, advertisement calls or displays, and pheromone production” (Fromhage et al. 2016). Several factors have been proposed to determine which of the two sexes contributes more towards mate searching (McCartney et al. 2012; Fromhage et al. 2016). However, sex differences in mate searching costs from predation risk, proposed as a potential determinant (Fromhage et al. 2016), has rarely been tested in natural populations (Heller 1992; Raghuram et al. 2015).

Sex differences in benefits of multiple matings, differential gametic and parental investments, and time- and predation-related costs have been proposed to determine which of the two sexes invests more effort in mate searching (Trivers 1972; Hammerstein and Parker 1987; Fromhage et al. 2016). Since males typically get higher maximum potential benefits from multiple matings than females (Bateman 1948; Rodríguez-Muñoz et al. 2010; Byers and Dunn 2012), and females are predicted to be the limiting resource to males by investing more in offspring (Andersson 1994), males are expected to be the searching sex. Contrary to this expectation, in a classic paper, Hammerstein and Parker (1987) showed that either sex can potentially search, arguing that sex differences in relative parental investment do not determine which sex searches for mates. But in most animals, males searching for females is the usual mode of pair formation (Darwin 1871; Thornhill 1979). Among other factors, the cost of mate searching has been suggested as a potential determinant of this asymmetry (Fromhage et al. 2016). Fromhage et al. (2016) proposed that higher costs for females could explain the prevalence of male searching, calling it the ‘sex-specific cost hypothesis’. One of the ubiquitous costs experienced during mate search is predation (Gwynne 1987; Magnhagen 1991; Heller and Arlettaz 1994; Zuk and Kolluru 1998; Zuk et al. 2006; Raghuram et al. 2015). Predation risk as a cost of mate searching for both sexes, however, has not been assessed.

Although mate searching by males is prevalent in most animals, in some other taxa such as orthoptera, females physically move to localise signalling males who are predominantly sedentary (Darwin 1871; Thornhill 1979). Thus, females share mate searching responsibilities, thereby exhibiting reduction in the asymmetry in mate searching between the sexes observed in many animal taxa. What factors explain females contributing towards pair formation and its maintenance? Thornhill (1979) attributes this to two potential factors: direct benefits provided by males to females on pair formation and/or the risks associated with signalling. In many species of orthoptera, on pair formation, males provide direct benefits such as a burrow for safe shelter (Gwynne 1995) or courtship nuptial gifts (Arnqvist and Nilsson 2000). McCartney et al. (2012) examined the interspecific mate searching differences across 32 taxa from a katydid genus, using theory and observational data. Their findings provide comparative evidence for the hypothesis that mate searching by females can be explained by the direct benefits offered by males if these are substantial. The alternative hypothesis states that since males benefit more from multiple mating, they should be selected to perform risky mate searching activities; hence, males are expected to face higher risks while signalling (Thornhill 1979; McCartney et al. 2012). The relative risk of signalling versus responding has however rarely been empirically tested.

Predation risk in the context of mate searching communication has predominantly been studied from the signaller’s perspective (Zuk and Kolluru 1998). Many studies have demonstrated how sexual advertisement in the form of conspicuous calls makes signallers vulnerable to predation (Walker 1964; Bell 1979; Ryan et al. 1982; Belwood and Morris 1987; reviewed in Zuk and Kolluru 1998). However, studies analysing predator diet found evidence for responding females being at an equal, if not higher predation risk, in comparison with signalling males (Heller and Arlettaz 1994; Raghuram et al. 2015). Thus, both signalling and responding to signals entail predation risk, since signalling attracts eavesdropping predators (Zuk and Kolluru 1998) and movement towards a signal increases exposure to predators (Gwynne 1987; Heller and Arlettaz 1994; Raghuram et al. 2015). There is however, a paucity of studies that attempt to estimate the relative predation risk of signallers and responders (but see Heller 1992; Raghuram et al. 2015).

We examined the mate searching costs from predation risk in tree crickets by estimating predation risk experienced by calling males and responding females in comparison with their non-communicating controls. For determining the intensity of selection due to predation on particular behaviours, the number of crickets captured by predators while exhibiting those behaviours of interest need to be quantified. We studied predation risk using an approach novel to the field of communication, though commonly used in ecology (Holling 1959; Lima and Dill 1990; Hebblewhite et al. 2005; Brechbühl et al. 2011). We defined the risk of predation “as the probability of being killed” while exhibiting the strategy of calling by males and responding to calls by females (Lima and Dill 1990). We partitioned risk into constituent parts, each characterised by a discrete spatial scale at which predator and prey interact: co-occurrence (spatial overlap from which predator and prey can potentially perceive each other), encounter (spatial proximity from which predators can potentially attack prey) and being eaten (the behavioural outcome once the predator attacks prey). At each scale, the binary response of prey either succeeding or failing to avoid the predator helped estimate probabilities. Predation risk was estimated as a product of these probabilities (represented in Fig. 1). Such a comprehensive approach is critical to determine any trade-offs across different scales that might not reflect in the total predation risk if studied only at the level of proportion of sexes in predator diets, predator visitations in playback experiments, or predator preferences of certain prey behaviours in a controlled environment (Tuttle and Ryan 1981, Heller and Arlettaz 1994, Alem et al. 2011). Furthermore, quantifying predator visits and/or predator preference towards calling males and responding females and equating it with intensity of selection assumes that both those encounters happen with the same frequency in the wild. Hence, we formally estimated the probability of these encounters in the field, in addition to the respective capture probabilities, and multiplied them in order to estimate predation risk.

**Fig. 1.**
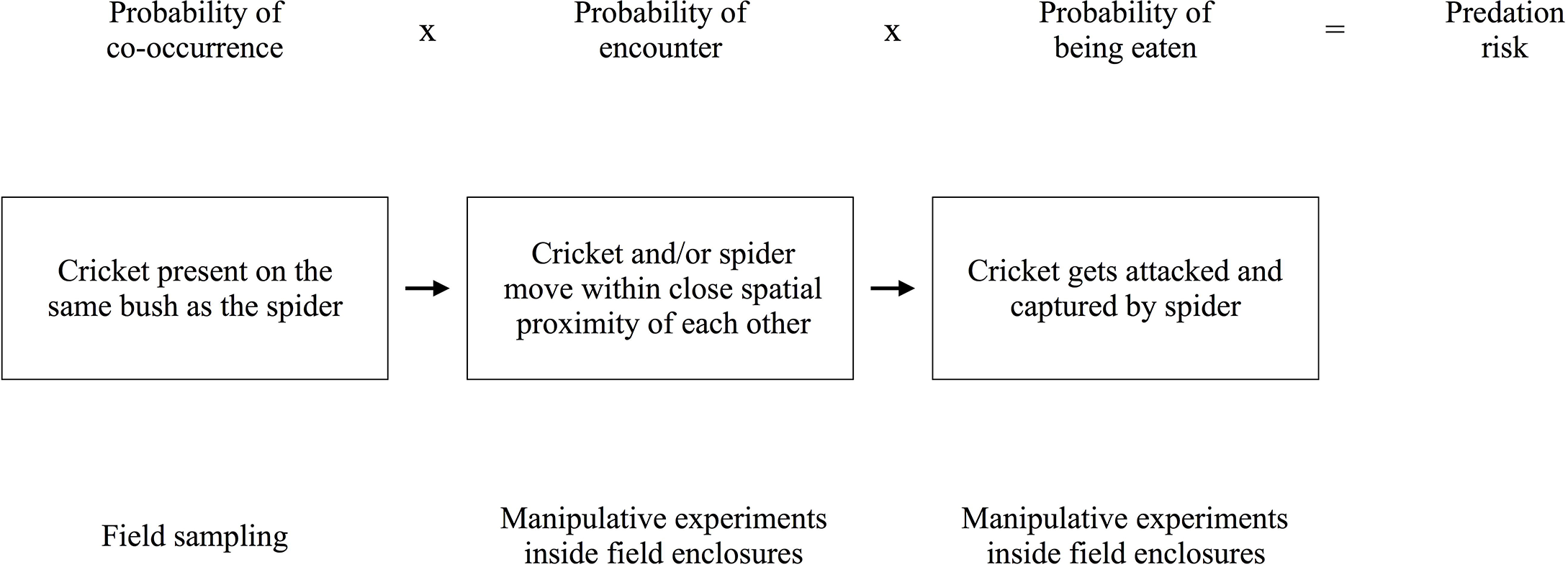
Graphical representation of experimental design employed in the study to estimate predation risk. Each box represents a cricket interacting with its predator, a spider, at a particular spatial scale. Each arrow represents transition to the next spatial scale. Text above each box indicates what constituent of predation risk is studied and text below indicates how it was estimated. Probabilities from each spatial scale were multiplied to estimate predation risk.

We estimated predation risk as a probability of mortality experienced by male tree crickets when they were calling and not calling, and female crickets when they were responding and not responding to calls, from their main predators, green lynx spiders to test two hypotheses. The sex-specific cost hypothesis attributes ubiquitous male mate searching in most animal taxa to higher costs of searching in females (Fromhage et al. 2016) and expects females to search less if mortality during searching is higher for females in comparison with the same for males. In a system where both males and females contribute towards mate searching, the sex-specific cost hypothesis expects mortality experienced while searching for mates to be similar in both males and females. The second hypothesis we tested expects searching activity by males to be more risky than that by females since males benefit more from multiple matings.

## Methods

We carried out our study on a tree cricket species, *Oecanthus henryi* whose main predator is the green lynx spider, *Peucetia viridans*. *Oecanthus henryi* is found extensively in the dry scrubland of southern India, predominantly on bushes of *Hyptis suaveolens*. *Oecanthus* species exhibit a mating system typical of true crickets (Gryllidae), where the males produce a long-range species-specific call and females do not call, but detect, recognize and locate males of their species (Walker 1957). Males of *O. henryi* typically call from *H. suaveolens* leaves and the females negotiate the complex architecture of the bushes and approach the calling males (Bhattacharya 2016). *Oecanthus henryi* male calls are made up of rhythmic chirps (Metrani and Balakrishnan 2005). *Peucetia viridans* (family Oxyopideae) is commonly observed on *Hyptis suaveolens* bushes and has been observed preying upon tree crickets and honeybees (VRT and RB, personal observations). Spiders perceive acoustic signals (Lohrey et al. 2009) as both airborne acoustic cues (Shamble et al. 2018) and substrate-borne vibratory cues (Barth 2002), the former most likely detected by the air-flow-sensitive hairs present in abundance on spider bodies. Similarly, spiders may perceive locomoting females using the vibratory cues produced by females while moving on bush branches.

All field surveys, sampling sessions and experiments were carried out in a homogenous patch of *H. suaveolens* on unused farmland, near Ullodu village (13°38’27.2”N 77°42’01.1”E) in Chikkaballapur district of Karnataka state in southern India. Laboratory experiments were performed on the campus of the Indian Institute of Science, Bangalore. All experiments in semi-natural conditions were carried out in an outdoor enclosure, constructed with a steel frame of dimensions 6m x 6m x 3m and fastened with a fibre mosquito mesh (mesh size: 0.1cm x 0.2 cm), on the campus of Indian Institute of Science, Bangalore.

We estimated predation risk as a product of three probabilities: 1) probability of co-occurrence of *O. henryi* and *P. viridans* on a bush (POC), 2) probability of *O. henryi* encountering *P. viridans*, given their co-occurrence (POE), and 3) probability of *O. henryi* being eaten by *P. viridans*, given an encounter (POBE).

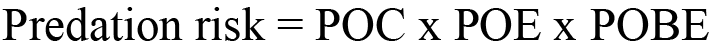

The study was carried out in four parts. First, we performed extensive sampling to investigate who the main predators of *O. henryi* were in the field. Second, we sampled a population of *O. henryi* in the field to estimate the probability with which they co-occur with their main predator species, considering a single *H. suaveolens* bush as a unit (POC). Then we conducted two experiments in a semi-natural outdoor enclosure to obtain the probability of encounter between *O. henryi* and *P. viridans* when both co-occurred on the same bush (POE), and the probability with which *O. henryi* are eaten by *P. viridans* upon encounter (POBE, represented in Fig. 1). The sampling and experiments to measure the probabilities were carried out on four different treatment regimes: calling males, non-calling males, phonotaxis performing females and non-responding females of *O. henryi*.

### Predator sampling

Not much is known about the identity of predators of *Oecanthus* species (Ponce-Wainer and Del Castillo 2008, but see O’Neill and O’Neill 2003, Ercit 2013). We conducted a series of experiments and sampling sessions to elucidate the predators of *O. henryi*. Playback experiments were carried out in the field to determine acoustically orienting aerial predators of *O. henryi*. Visitations by aerial predators were compared at speakers playing back *O. henryi* calls with paired silent speakers for a total of 40 hours (experimental details in supplementary information section S 1). For discovering arboreal predators of *O. henryi*, relative abundance sampling sessions and predation experiments were carried out. The extensive relative abundance sampling helped determine who the potential predators of *O. henryi* were (sampling details in supplementary information section S 2.1). To establish whether these potential predators were real predators, detailed predation experiments were conducted in the field (details in supplementary information section S 2.2 and S 2.3). Experiments were also conducted to understand how starvation period of the predator affects predation so as to better design further experiments (experimental details in supplementary information section S 2.4).

### Probability of co-occurrence

To determine the probability of co-occurrence between *O. henryi* and its main predator *P. viridans*, incidences of *O. henryi* co-occurring with large *P. viridans* (body size larger than 5.12 mm; for details refer to results section and supplementary information section S 2.2) on the same bush were recorded in the field between November and May. Between 1900hrs and 2115hrs, calling males were located using their calls, whereas non-calling males and females were localized using 5×5 m quadrat sampling or opportunistically. These plots were chosen by dividing the whole field into 5×5 m plots and randomly selecting from them. The field was made up mostly of *Hyptis suaveolens* bushes. Once a quadrat was chosen, all bushes in it were sampled for the presence of non-calling males and females. Once localized, both male and female crickets, were focally sampled for at least 30 minutes. This time was allocated mainly for males to call, to help distinguish callers from non-callers. The bush or bushes, on which these crickets were observed while sampling, were thoroughly searched for the presence of *P. viridans* at the end of the sampling session. If a spider was present, distances between the cricket and the spider and whether they were present on the same or different branches was recorded, along with the height and distance of the spider from the centre of the bush. Post sampling, crickets and spiders were collected and brought back to the field station for marking and sizing, respectively. Crickets were marked with a unique tricolour code using nontoxic paint markers (Edding 780, Edding, St Albans, U.K.), to avoid resampling; spiders were sized to confirm their ability to predate on crickets (body size larger than 5.12 mm; for details refer to results section and supplementary information section S 2.2), and both were released back into the field. Male *O. henryi* were considered as callers based on whether or not they called more than 20% of the time they were observed whereas a male that did not call at all was considered as a non-calling male. 20% of the calling effort was chosen as a cut-off so as to avoid choosing infrequent signallers as callers. To be certain that this decision does not affect our results, we ran all the analyses with a threshold of both 10% and 30% of the calling effort for selecting calling males and found that the results do not change qualitatively. Since there was no intuitive way to categorise communicating and non-communicating females during field observations, observed females were randomly classified as responding and non-responding based on a supplementary experiment (that estimated the relative frequency of communicating, or phonotactic, females in the same population). This random sampling involved computationally segregating (sampled for 10,000 iterations) the observed females into responding and non-responding females, using the proportion of wild-caught females known to be responsive (0.3; details in supplementary information section S 3).

### Probability of encounter given co-occurrence

To determine the probability of encounter between *O. henryi* and *P. viridans* given their co-occurrence on a bush, experiments were carried out in semi-natural conditions, inside a large outdoor enclosure. Crickets and spiders were collected from wild populations from in and around Peresandra, Karnataka, India (13°35’25.3”N 77°46’50.4”E), a few days before the experiment. Crickets were maintained on *H. suaveolens* bushes and spiders were provided *Gryllus bimaculatus* nymphs 2-3 times a week, both in the laboratory. Female crickets were collected as nymphs from the field and fed on an apple diet till they eclosed into adults after which they were maintained on *H. suaveolens* bushes. This exercise ensured virginity of all tested females, which increases the propensity of females to perform phonotaxis (Modak, Brown and Balakrishnan, unpublished results). Male and female crickets were maintained separately. Spiders were maintained in individual plastic boxes (6 cm diameter, 4 cm height). Spiders were starved for 48 hours prior to a trial and crickets were transferred to *H. suaveolens* bushes in outdoor cages at least a day before a trial to acclimatise them. No cricket was repeated across or within treatments and no spiders were repeated within treatments.

Each trial involved releasing one cricket and one spider on a *H. suaveolens* bush. Crickets were released on bushes at least 4 hours before the trial was started. The spiders were released at 1900 hours, marking the beginning of a trial. They were released on the bush at a height and distance from its centre, picked randomly from the interquartile range of their respective uniform distributions that were obtained from field data (height range on the bush for male crickets: 29 to 51 cm; for female crickets: 27 to 56 cm; for spiders on bushes with male crickets: 36 to 67 cm; for spiders on bushes with female crickets: 19 to 50 cm; Distance from centre of bush for male crickets: 12 to 24 cm; for female crickets: 11 to 29 cm; for spiders on bushes with male crickets: 11 to 22 cm; for spiders on bushes with female crickets: 12 to 21 cm). Spiders were released on either the same or on a different branch as the cricket at 1900 hours with the proportion with which they were observed in the field (0.154 of all co-occurrences between crickets and spiders in the field were on the same branch for males, and 0.286 for females. These different proportions for males and females were not significantly different). On the same or different branch, the distance at which spiders were released from crickets was drawn randomly from the interquartile range of a uniform distribution of distances at which crickets and spiders were observed in the field (distance between spider and male cricket on same branch: 7 to 9 cm; on different branch: 23 to 40 cm; distance between spider and female cricket on same branch: 10 to 15 cm; on different branch: 19 to 38 cm). Once the spider was released at 1900 hours, the interaction between the cricket and spider was observed for about 120 minutes by scan sampling each individual alternately every 30 seconds. An encounter was defined as any spatial proximity between the cricket and spider, within 4 cm of each other, on the same branch, that includes spiders capturing crickets, spiders unsuccessfully attacking crickets or either spider or cricket moving away without spiders attacking (Table S1 in supplementary information). This distance was the outer range from which *P. viridans* attacked *O. henryi* and also the distance at which the cricket could potentially antennate the spider (VRT, personal observations).

The 4 treatments considered for this experiment were, calling males, non-calling males, phonotaxis performing females and non-phonotactic females. For the responding female treatment, female crickets were played back conspecific male calls from a speaker (X-mini Capsule Speaker V1.1, Xmi Pte Ltd, Singapore) placed 60 cm away, across the bush from the female’s position, at 1900 hrs. The observation was considered a trial only if the female performed phonotaxis and reached within 20 cm of the speaker. The speaker was fixed on a stand, which was adjusted to the median height at which calling males were observed in the field (42 cm). The SPL of each call broadcast from the speaker was adjusted to be 61 dB (r.m.s. re. 2 x 10^-5^ Nm^-2^) at the female’s location with the help of a ½” microphone (Brüel and Kjær A/S, Denmark, Type 4189, 20 Hz to 20 kHz) fitted on a Sound Level Meter Type 2250 (Brüel and Kjær A/S, Denmark). Since the call carrier frequency changes with temperature in *O. henryi*, the choice of call to be played back was based on the temperature recorded at 1900 hours every evening. The call was chosen from among 3 representative calls that were recorded at 22°C, 24°C and 26°C (calls recorded by Rittik Deb, Deb 2015), whichever was closest to the recorded temperature. This call was played back in a loop using Audition software (Adobe, Version 5.0.2) on a MacBook Pro (2011) laptop using X-mini (Capsule Speaker V1.1, Xmi Pte Ltd, Singapore) speakers, for the entire duration of the experiment. For the non-responding female treatment, the same setup as above was maintained but no call was played back.

### Probability of being eaten given encounter

The probability of *O. henryi* being eaten by *P. viridans* once encounter occurs was examined empirically in the same outdoor semi-natural setup. *O. henryi* and *P. viridans* were collected and maintained using the same protocol mentioned in the earlier section. The same four treatments were maintained for this experiment. Crickets were released at least 4 hours before the commencement of the experiment at 1900 hours on the bush at a height and distance from its centre as explained in the last experiment. From 1900 hours onwards, focal observations were made for at least 45 minutes to allow males to call and to allow females to perform phonotaxis. Following these observations and based on whether the male called and female performed phonotaxis, a spider was gently released within close proximity of the cricket, not more than 6 cm away from it, using a *H. suaveolens* stick. A trial was considered only if the spider attacked, which was confirmed by videotaping all interactions. A spider capturing the cricket was scored as the cricket being eaten by the spider.

### Analyses

Since comparing estimates of rare and non-normally distributed events can be challenging, we employed non-parametric bootstrapping and permutation tests (Manly 2006). These are robust methods for obtaining confidence intervals and *P*-values, respectively, since they make few assumptions about the underlying distributions (Manly 2006; Nakagawa and Cuthill 2007). Bootstrapping was used to generate 95% confidence intervals around each probability. This process involved sampling with replacement, for 10,000 iterations, from the original vector of success/ failures used to calculate the probability. Overlap in confidence intervals was used to infer statistical significance for each relevant comparison (Cumming and Finch 2005). Additionally, permutation tests were carried out to assess statistical significance (Manly 2006). We used software R, Version 3.3.3 (R Core Team 2017) to run all analyses, and the ‘ggplot2’ package (Wickham 2009) to plot all graphics.

## Results

### Predator sampling

Playback experiments were carried out in the field to observe if there are any acoustically orienting aerial predators of *O. henryi*. Bats (species unknown) flew past the playback speaker on 4 separate occasions, and past the control speaker on 5 occasions out of a total of 40 hours of observation. Bats flew past both speakers in similar numbers. Also, a praying mantis approached a broadcast speaker once. In relative abundance sampling to investigate arboreal predators, 15 5×5m plots were sampled, amounting to 1083 bushes. A total of 127 *P. viridans* individuals, 129 spiders belonging to the web-building guild, and 1 praying mantis were observed, along with many beetles, roaches, termites and moths which were not enumerated since they are not potential predators of tree crickets. Of these, *P. viridans* and spiders belonging to the web-building guild were categorized as potential predators. In the field predation experiment, 16 out of 30 *P. viridans* that were offered *O. henryi*, captured and consumed them. Mean size of *P. viridans* that successfully predated on *O. henryi* was 9.12 mm (n = 16) and the mean size of those that did not predate was 4.22 mm (n = 14), and they were significantly different (Randomisation test, *P* < 0.001). All *P. viridans* that captured *O. henryi* were larger than 5.12 mm in body length (n = 16) (Fig. 2). In similar sets of experiments, spiders belonging to the web-building guild, were found not to be main predators of *O. henryi* (details in supplementary information section 2.3).

**Fig. 2.**
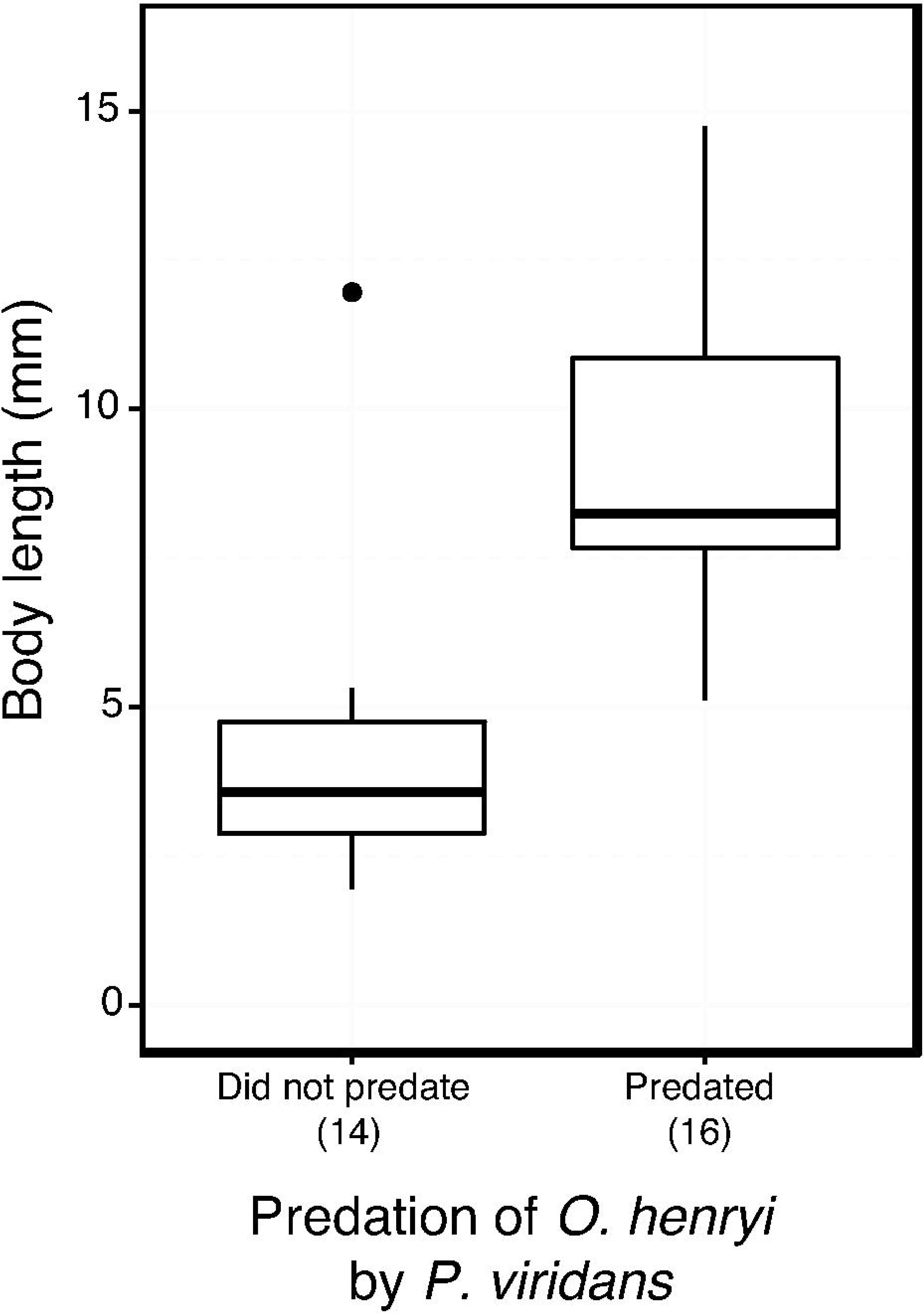
Size distinction between *P. viridans* individuals that predated on *O. henryi* and the ones that did not. All spiders that captured crickets were larger than 5.12mm in body length.

### Probability of co-occurrence

The probability with which crickets co-occur with spiders was similar between calling males, non-calling males, responding females and non-responding females. Pairwise comparisons between calling males and non-calling males (*p* = 0.07), and between calling males and responding females (*p* = 0.78) reveal no statistically significant differences (Fig. 3a). These results did not qualitatively change when all sampled females (n = 43) were considered instead of only the randomly sampled segregate classified as responding females (*p* = 0.79). Since responding females and non-responding females were categorised by randomly assigning females to these two groups, statistical differences between the two groups can be attributed to Type 1 error. Hence, the statistical hypothesis test for a difference between responding and non-responding females has not been presented (Fig. 3a).

**Fig. 3.**
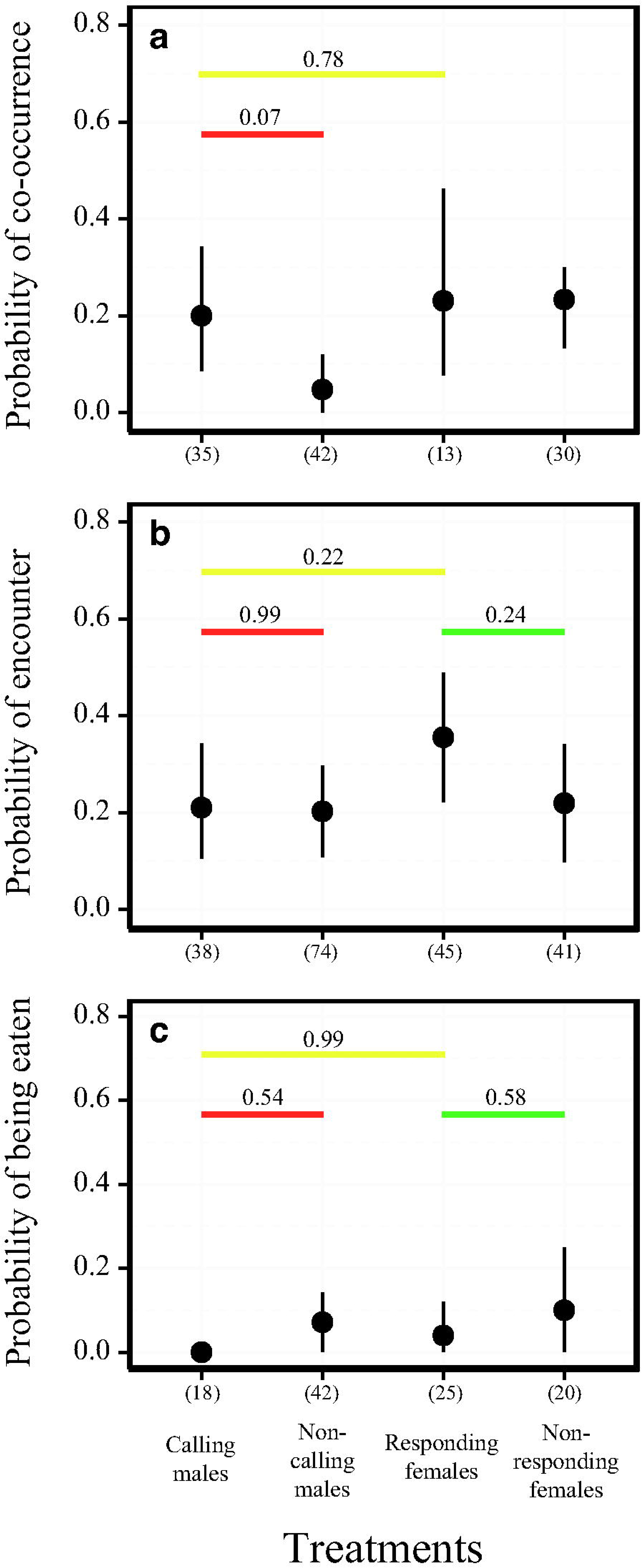
Constituent probabilities of predation risk faced by *O. henryi* from its predator, *P. viridans* partitioned into three spatial scales: probability of (a) co-occurrence, (b) encounter and (c) being eaten. The different treatments of *O. henryi* include calling males, non-calling males, responding and non-responding females. Bootstrapped probabilities are shown along with 95% confidence intervals and sample sizes are mentioned in parentheses. Pairwise comparisons between calling and non-calling males (red dashes), responding and non-responding females (green dashes), and calling males and responding females (yellow dashes) are represented with associated *p* values.

### Probability of encounter given co-occurrence

Once co-occurring on the same bush, crickets encounter spiders with similar probabilities. None of the pairwise comparisons between calling males and non-calling males (*p* = 0.99), responding females and non-responding females (*p* = 0.24), and between calling males and responding females (*p* = 0.22) were significantly different from each other (Fig. 3b).

### Probability of being eaten given encounter

When an encounter was forced between the spiders and crickets, relatively few crickets, regardless of sex and activity were captured and eaten by *P. viridans* (Fig. 3c). Thus, on encounter, spiders capture and eat crickets across the 4 treatments with similar probability. None of the pairwise comparisons between the probabilities of being eaten of calling males and non-calling males (*p* = 0.54), responding females and non-responding females (*p* = 0.58), and between calling males and responding females (*p* = 0.99) were significantly different from each other (Fig. 3c). The probability of being eaten is much lower in this experiment compared to the field predation experiment performed to establish *P. viridans* as the real predator (supplementary information section S 2.2) because the field predation experiment was conducted inside plastic boxes, in a restricted space, where crickets could not escape, unlike when on the bush.

### Predation risk

The resulting product of these probabilities (product of co-occurrence, encounter and capture probabilities, Fig. 1), the predation risk, is also similar between all 4 treatments. When compared in a pairwise manner, calling males and non-calling males (*p* = 0.36), responding females and non-responding females (*p* = 0.83), and between calling males and responding females (*p* = 0.56), were not significantly different from each other (Fig. 4). We suspected the predation risk for all four treatments to be similar mainly because of the similar probabilities of being eaten by spiders (Fig. 3c). Hence, we measured the product of the probability of co-occurrence and probability of encounter, without including the probability of being eaten. The lack of differences persist in the pairwise comparisons of calling males and non-calling males (*p* = 0.59), responding females and non-responding females (*p* = 0.77), and between calling males and responding females (*p* = 0.52) (supplementary information Fig. S2).

**Fig. 4.**
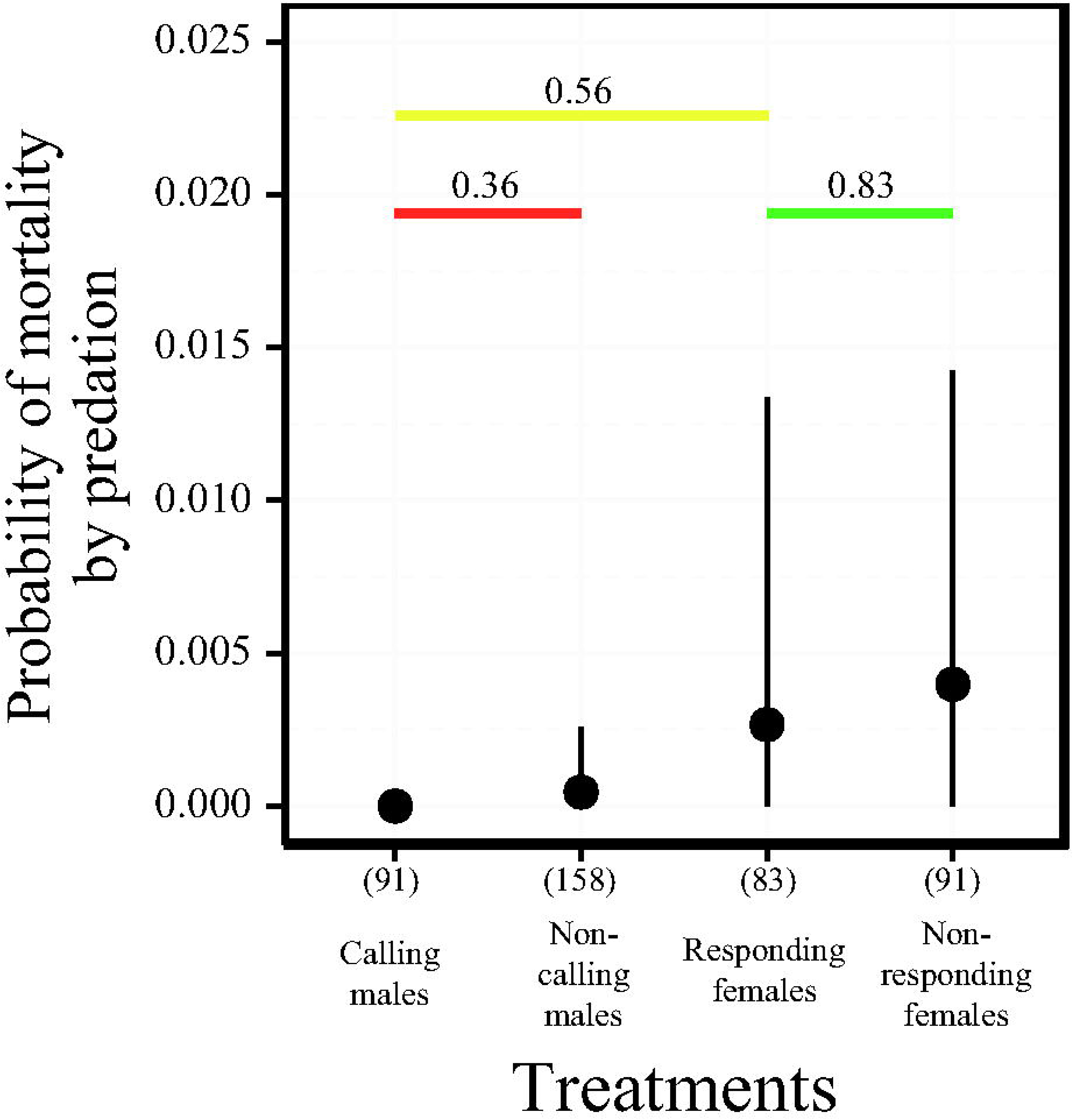
Predation risk faced by *O. henryi* from its predator, *P. viridans*. The different treatments of *O. henryi* include calling males, non-calling males, responding and non-responding females. Probabilities are bootstrapped values represented as 95% confidence intervals. Values in parentheses are sample sizes. Pairwise comparisons between calling and non-calling males (red dashes), responding and non-responding females (green dashes), and calling males and responding females (yellow dashes) are represented with associated *p* values.

## Discussion

Our study investigated the relative predation risk of communicating individuals, signallers and responders, in natural populations on a within-night time scale. Observing interactions at various spatial scales allowed a comprehensive quantification of predation risk by taking into account predator-prey dynamics. Our choice of predator was justified using extensive field surveys and acoustic sampling sessions to determine all potential predators and carefully pruning that list using predation experiments to select a predator that is ecologically relevant to our system of choice. We found that the green lynx spider *P. viridans* was the main predator of *O. henryi*. Spiders have been observed to be predators of several species of crickets (Hedrick and Kortet 2006; Dangles et al. 2006; Storm and Lima 2010). Spiders are sensitive to both long-range acoustic (Shamble et al. 2016) and substrate-borne vibratory cues (Barth 2002). Such multimodal sensitivity could allow them to perceive both acoustic calls and locomotory cues, making spiders good model predators for studying costs of communication that involves calling by males, and movement by females.

Predation risk of calling and non-calling males and females responding and not responding to calls was similar across all spatial scales relevant to predator-prey dynamics. We examined and compared the probability of communicating and non-communicating male and female crickets co-occurring on bushes with spiders. This probability was similar, indicating that risk faced from distribution of spiders across bushes is not influenced by whether the cricket is a communicating or non-communicating male or female. These females were classified as responding females based on random sampling of wild females based on the limited information we had, and a future step would be to compare the co-occurrence patterns of communicating and non-communicating females. Based on the smaller sample size of wild females tested for motivation, anecdotal observations suggest that the co-occurrence of communicating and non-communicating individuals is not very different. While co-occurring on the same bush, the probability of encountering a spider was similar between the predominantly sedentary calling males when compared with mostly mobile responding females and non-calling males. Similarly, encounter probability with a spider was similar between mobile responding females when compared with non-responding females and calling males. A possible explanation for this result is that communicating and non-communicating individuals are taking similar necessary evasive measures to spatially avoid predators (reviewed in Sih 2005). It will be interesting to investigate whether the two sexes spatially avoid predators. In addition, we investigated the probability with which crickets were captured by spiders when attacked and found no differences. This probability was extremely low for both sexes in comparison with the earlier two probabilities. This result is not unexpected since crickets are known to have high escape success against spider attacks, attributable to their air-sensing systems (Dangles et al. 2006). Although the probability of calling males being eaten by spiders was zero, it was statistically similar to the non-zero probability of non-calling males and responding females. Finally, the resultant product of the three probabilities, considered as predation risk, was extremely low for the two sexes, primarily due to high escape probabilities exhibited by crickets when attacked by spiders. Also, this probability was similar for all four treatments, suggesting the cost from predation was comparable for communicating and non-communicating male and female crickets. Categorising predation risk according to not only sex, but also behaviour, allowed us to estimate and interpret the predation risk experienced by the two sexes when they were searching for mates and when they were not. Since predation risk between communicating individuals was found to be similar, we can infer that the two different mate searching strategies, signalling and moving, carry equivalent, low levels of risk. Similarly, since risk from predation was similar between communicating and non-communicating individuals, the cricket’s choice to search for mates does not expose it to greater predation risk.

The sex-specific cost hypothesis predicts sex differences in mate searching effort if searching is more costly for females than for males. Since in orthopterans both sexes share mate searching effort, we expected costs from predation risk between the sexes to be more similar. Our results, that costs of mate searching from predation risk are similar for signalling males and responding females, support the sex-specific cost hypothesis as an explanation of shared mate searching responsibilities in orthopterans. Furthermore, since there were no differences between communicating and non-communicating individuals, the predation risk associated with mate searching is nearly zero. More such examinations of mate searching costs between the sexes are important to determine whether searching costs indeed are systematically higher for females than males. Furthermore, it will be interesting to test whether the extent of sexual asymmetry in mate searching corresponds to the differences in sex-specific costs, across different taxa. This should include mating systems ranging from only male searching to only female searching, in a comparative framework.

We also tested the predictions of one of the two factors proposed to explain females sharing mate searching responsibilities with males in long distance signalling taxa: high risk of signalling (Thornhill 1979). Risk associated with signalling has been shown in several taxa in which males signal for mate searching (Walker 1964, Bell 1979, Ryan et al. 1982, Belwood and Morris 1987, reviewed in Zuk and Kolluru 1998). We suggest that studies testing this hypothesis should not only provide evidence that the risk is high for signalling males, but also that this risk is higher than that for responding females. Very few studies have estimated risk of signalling in comparison with searching. Our findings, in corroboration with two other studies (Heller 1992, Raghuram et al. 2014), are not consistent with the prediction that males are selected to perform the more risky mate searching activity. The hypothesis that direct benefits offered by males drives evolution of female mate searching was however supported by theoretical and observational results in katydids (McCartney et al. 2012). In addition, courtship feeding as observed in *Oecanthus* species (Houghton 1909; Fulton 1915), has been shown to be an ancestral trait in the orthopteran suborder Ensifera, suggesting that males of most Ensiferan genera will offer direct benefits to their female counterparts on pair formation (Gwynne 1997). Furthermore, direct benefits offered by males might offset the costs females experience while responding to signals that are generated by sedentary males. Since males are expected to benefit from multiple matings and females from direct benefits, the costs can also be expected to be shared between the sexes, a potential outcome supported by our results. Hence, females contributing towards mate searching efforts in orthopterans can perhaps be better explained by provision of resources by males to females rather than invoking higher risk of signalling.

In conclusion, although predation risk of signalling in males has been considered to be high (Zuk and Kolluru 1998), when compared with risk of responding, our findings show these risks to be similar. It is only by addressing predation risk between the communicating sexes across several relevant spatial scales that we could compare risks faced by the two mate searching strategies. More comparative studies on different species on predation associated costs between the mate searching sexes would help update our understanding of whether at all there are systematic cost differences in mate searching strategies. Finally, we also show that overall predation costs of communication per night are low and that a predation event is very rare, which raises questions on the importance of predation as a major selection pressure on the evolution of communication.

## Author contributions

VRT participated in conceptualising and designing the study, carried out data collection and analysis and wrote the manuscript. KI contributed to data analysis, interpretation and writing the manuscript. RB contributed to conceptualising and designing the study, interpretation of data and writing the manuscript.

## Acknowledgements

We thank the DBT-IISc partnership programme (2012-2017) of the Department of Biotechnology, Government of India for funding the fieldwork and Ministry of Human Resource Development, Government of India for funding the research fellowship of VRT. We thank the DST-FIST program of the Govt. of India for some of the equipment used in the study. We thank Diptarup Nandi, Rittik Deb and Monisha Bhattacharya for helpful discussions and Diptarup Nandi for comments on the manuscript. We thank Mr. B. Sridhar (Garden and Nursery Technical Officer) and Nursery staff for their support. We would like to thank Ayan, Babu, Chaithra, Chirag, Dakshin, Girish, Himamshu, Harsha, Ismail, Lakshmipriya, Manoj, Meenakshi, Meghana, Murthy, Shivaraju, Sidanth, Sonu, Sriniketh, Srinivasan, Sunaina and Vinayaka for their help in making behavioural observations and Manjunatha Reddy for his assistance in fieldwork.

## Conflict of Interest

The authors declare that they have no conflict of interest.

## Data Availability statement

The datasets generated and analysed during the current study can be accessed at https://doi.org/10.6084/m9.figshare.7794131.v1

## Ethics statement

All behavioural data sampling and experiments were performed in accordance with the national guidelines for the ethical treatment of animals laid out by the National Biodiversity Authority (Government of India).

## References

Alem S, Koselj K, Siemers BM, Greenfield MD (2011) Bat predation and the evolution of leks in acoustic moths. Behav Ecol Sociobiol 65:2105–2116

Andersson MB (1994) Sexual selection, Princeton University Press, Princeton

Arnqvist G, Nilsson T (2000) The evolution of polyandry: multiple mating and female fitness in insects. Anim Behav 60:145–164

Balakrishnan R (2016) Behavioral ecology of insect acoustic communication. In: Pollack GS, Mason AC, Popper AN, Fay RR (eds) Insect hearing. Springer International Publishing, Switzerland, pp 49–80

Barth FG, (2002) A spider’s world: senses and behavior. Springer-Verlag, New York

Bateman AJ (1948) Intra-sexual selection in Drosophila. Heredity 2:349–368

Bell PD (1979) Acoustic attraction of herons by crickets. J N Y Entomol Soc 87:126– 127

Belwood JJ, Morris GK (1987) Bat predation and its influence on calling behavior in neotropical katydids. Science 238:64–67

Bhattacharya M (2016) Investigating pattern recognition and bi-coordinate sound localization in the tree cricket species. PhD dissertation, Centre for Ecological Sciences, Indian Institute of Science, Bangalore, India.

Brechbühl R, Casas J, Bacher S (2011) Diet choice of a predator in the wild: overabundance of prey and missed opportunities along the prey capture sequence. Ecosphere 2:1–15

Byers J, Dunn S (2012) Bateman in nature: predation on offspring reduces the potential for sexual selection. Science 338:802–804

Core Team R (2017) A language and environment for statistical computing. R Foundation for Statistical Computing. http://www.R-project.org.

Cumming G, Finch S (2005) Inference by eye: confidence intervals and how to read pictures of data. Am Psychol 60:170–180

Darwin C (1871) The descent of man and selection in relation to sex. John Murray, London, UK

Dangles O, Ory N, Steinmann T, Christides JP, Casas J (2006) Spider’s attack versus cricket’s escape: velocity modes determine success. Anim Behav 72:603–610

Deb R (2015) Mate choice, mate sampling and baffling behaviour in the tree cricket. PhD dissertation, Centre for Ecological Sciences, Indian Institute of Science, Bangalore, India.

Ercit K (2013) Size and sex of cricket prey predict capture by a sphecid wasp. Ecol Entomol 39:195–202

Fromhage L, Jennions MD, Kokko H (2016) The evolution of sex roles in mate searching. Evolution 70:617–624

Fulton BB (1915) The tree crickets of New York: life history and bionomics. N Y Agric Exp Station Tech Bull 42:1–47

Gwynne DT (1987) Sex-biased predation and the risky mate-locating behaviour of male tick-tock cicadas (Homoptera: Cicadidae). Anim Behav 35:571–576

Gwynne DT (1995) Phylogeny of the Ensifera (Orthoptera): A hypothesis supporting multiple origins of acoustical signalling, complex spermatophores and maternal care in crickets, katydids, and weta. J Orthop Res 4:203–218

Gwynne DT (1997) The evolution of edible ’sperm sacs’ and other forms of courtship feeding in crickets, katydids and their kin (Orthoptera: Ensifera). In: Choe JC and Crespi BJ (eds) The evolution of mating systems in insects and arachnids. Cambridge University Press, Cambridge, UK, pp 110–129

Hammerstein P, Parker GA (1987) Sexual selection: games between the sexes. In: Bradbury JW, Andersson MB (eds) Sexual selection: testing the alternatives. John Wiley & Sons, Chichester, UK, pp 119–142

Hebblewhite M, Merrill EH, McDonald TL (2005) Spatial decomposition of predation risk using resource selection functions: an example in a wolf–elk predator–prey system. Oikos 111:101–111

Hedrick AV, Kortet R (2006) Hiding behaviour in two cricket populations that differ in predation pressure. Anim Behav 72: 1111–1118

Heller KG (1992) Risk shift between males and females in the pair-forming behavior of bushcrickets. Naturwissenschaften 79:89–91

Heller KG, Arlettaz R (1994) Is there a sex ratio bias in the bushcricket prey of the scops owl due to predation on calling males? J Orthop Res 2:41–42

Holling CS (1959) The components of predation as revealed by a study of small-mammal predation of the European Pine Sawfly. Can Entomol 91:293–320

Houghton CO (1909) Observations on the mating habits of Oecanthus. Entomol News, 274–279

Lima SL, Dill LM (1990) Behavioral decisions made under the risk of predation: a review and prospectus. Can J Zool 68:619–640

Lohrey AK, Clark DL, Gordon SD, Uetz GW (2009). Antipredator responses of wolf spiders (Araneae: Lycosidae) to sensory cues representing an avian predator. Anim Behav 77:813–821

Magnhagen C (1991) Predation risk as a cost of reproduction. TREE 6:183–186.

Manly BFJ (2006) Randomization, Bootstrap and Monte Carlo Methods in Biology, 3rd edn. Chapman and Hall/CRC, New York

McCartney J, Kokko H, Heller KG, Gwynne DT (2012). The evolution of sex differences in mate searching when females benefit: new theory and a comparative test. - Proc R Soc Lond B Biol Sci 279:1225–1232

Metrani S, Balakrishnan R (2005) The utility of song and morphological characters in delineating species boundaries among sympatric tree crickets of the genus Oecanthus (Orthoptera: Gryllidae: Oecanthinae): a numerical taxonomic approach. J Orthop Res 14:1–16

Nakagawa S, Cuthill IC (2007) Effect size, confidence interval and statistical significance: a practical guide for biologists. Biol Rev Camb Philos Soc 82:591– 605

O’Neill KM, O’Neill RP (2003) Sex allocation, nests, and prey in the grass-carrying wasp Isodontia mexicana (Saussure) (Hymenoptera: Sphecidae) - J Kans Entomol Soc 76:447–454

Ponce-Wainer JX, Del Castillo RC (2008) Female mate choice and no detected predation risk in relation to the calling song of Oecanthus niveus (Gryllidae: Oecanthinae). Ann Entomol Soc Am 101:260–265

Raghuram H, Deb R, Nandi D, Balakrishnan R (2015) Silent katydid females are at higher risk of bat predation than acoustically signalling katydid males. Proc R Soc Lond B Biol Sci 282:20142319

Rodríguez-Muñoz R, Bretman A, Slate J, Walling CA, Tregenza T (2010) Natural and sexual selection in a wild insect population. Science 328:1269–1272

Ryan MJ, Tuttle MD, Rand SA (1982) Bat predation and sexual advertisement in a neotropical anuran. Am Nat 119:136–139

Shamble PS, Menda G, Golden JR, Nitzany EI, Walden K, Beatus T, Elias DO, Cohen I, Miles RN, Hoy RR (2016) Airborne acoustic perception by a jumping spider. Curr Biol 26:2913–2920

Sih A (2005) Predator-prey space use as an emergent outcome of a behavioral response race. In: Barbosa P, Castellanos I (eds) Ecology of predator-prey interactions. Oxford University Press, New York, pp 240–255

Storm JJ, Lima SL (2010) Mothers forewarn offspring about predators: a transgenerational maternal effect on behavior. Am Nat 175:382–390

Tuttle MD, Ryan MJ (1981) Bat predation and the evolution of frog vocalizations in the neotropics. Science 214:677–678

Thornhill R (1979) Male and female sexual selection and the evolution of mating strategies in insects. In: Blum M, Blum N (eds) Sexual selection and reproductive competition in insects, Academic Press, New York, pp 81–121

Trivers RL (1972) Sexual selection and parental investment. In: Campbell BG (ed) Sexual selection and the descent of man, Aldine Press, Chicago, pp 136–179

Walker TJJ (1957) Specificity in the response of female tree crickets (Orthoptera, Gryllidae, Oecanthinae) to calling songs of the males. Ann Entomol Soc Am 50: 626–636

Walker TJJ (1964) Experimental demonstration of a cat locating orthopteran prey by the prey’s calling song. The Florida Entomol 47:163–165

Wickham W (2009) ggplot2: Elegant graphics for data analysis. Springer-Verlag, New York

Zuk M, Kolluru GR (1998) Exploitation of sexual signals by predators and parasitoids. Q Rev Biol 73:415–438

Zuk M, Rotenberry JT, Tinghitella RM (2006) Silent night: adaptive disappearance of a sexual signal in a parasitized population of field crickets. Biol Lett 2:521–524

