## Supplementary information for "Is the predation risk of mate-searching different between the sexes?"

**S 1 Aerial predator sampling**

*S 1.1 Experimental design details*

5x5m plots were randomly selected and a representative pre-recorded *O. henryi* male call was played back from close to a *H. suaveolens* bush in the middle of each plot (calls recorded by Rittik Deb, Deb 2015). The choice of call to be played back was based on the temperature measured that evening, and the closest matching temperature at which the calls were recorded. The sound pressure level (SPL) at which calls were played back was randomly drawn from a normal distribution of SPLs at which *O. henryi* males call (63.7±2.4 dB, Deb 2015). Speakers were placed at median male calling height above the ground (42 cm). This median male calling height was measured post focal sampling of calling males during the field observations to measure probability of co-occurrence (n = 35, Mean ± SD = 42.25 ± 16.48 cm). After calling males were observed for at least 30 minutes, height from the ground of the crickets’ last location were recorded, before they were collected. The randomly selected plot had a paired plot, selected randomly from any one of the 8 adjacent plots, in which a control experiment was simultaneously run. The control speaker was switched off during the trial. Focal observations were carried out for 2 hours every evening starting at 1900hrs, in two sets of experiment and control speakers, amounting to a total of 40 hours each at playback speakers and control speakers over 10 nights. Any visitation from a potential aerial predator was recorded if it flew within half a meter of the speaker (for results, refer to manuscript results section: Predator sampling).

**S 2 Arboreal predator sampling**

*S 2.1 Sampling details for determining the potential predators of O. henryi*

Since the field we worked in was too large to sample for absolute abundances of potential predators, relative abundance sampling – performed for determining who the potential predators were – was carried out in 15 5x5 m plots. These plots were chosen by dividing the whole field into 5x5 m plots and randomly selecting from them. All bushes above 20 cm were sampled for the presence of any potential predators, since bushes smaller than 20 cm barely had branches and leaves for potential predators to perch on. A total of 1083 bushes were observed carefully. Sampling included recording the heights and widths of all bushes observed, the potential predators observed and the height and distance from the centre of the bush at which they were observed. Only two kinds of predators, spiders belonging to the web-building guild and free-ranging spiders that do not build webs, *P. viridans* were abundant enough to be considered as potential predators (for results, refer to manuscript results section: Predator sampling).

*S 2.2 Experimental design details for determining whether P. viridans spiders are real predators of O. henryi*

To determine if *P. viridans* are real predators of *O. henryi*, a series of predation experiments in the field were performed. *O. henryi* was offered as prey to these potential predators to check if they do predate on the tree cricket species. Predators were sampled in randomly selected sub-fields of the Ullodu field and offered pre-collected *O. henryi* males under conditions where escape was difficult for the crickets, to test the predation capability of the potential predators. *P. viridans* were offered crickets inside plastic boxes (6 cm diameter, 4 cm height) (n = 30). This enclosed interaction was observed via scan sampling every 15 minutes either till the predator captured the cricket or at least 75 minutes. Body size of all *P. viridans* was recorded post the predation experiment, by measuring the length between anterior end of the cephalothorax and posterior end of the abdomen. All collected spiders were photographed along with a scale using a Nikon D40 digital camera, and the size was measured using ImageJ software (version 10.2) (for results, refer to manuscript results section: Predator sampling).

*S 2.3 Determining whether web-building spiders are real predators of O. henryi*

Once it was established that spiders belonging to the web-building guild are potential predators of *O. henryi*, predation experiments were carried out in the field to determine if they are real predators of *O. henryi*.

*S 2.3.1 Methods*

Web-building spiders of web diameter larger than 2 cm were localised in randomly selected sites in the field (n=47). This size threshold was chosen because webs smaller than diameters of 2 cm were too small to perform the experiment with. Once a spider was localized on its web, a predator-prey encounter was simulated by inducing *O. henryi* males to jump onto its web. Details of the encounter including whether the spider was able to capture the cricket, were noted. Only spiders with webs larger than a certain threshold diameter were observed to be capable of capturing *O. henryi*. To determine the abundances of web building spiders with web diameter larger than this threshold, another round of relative abundance sampling was carried out in the study site and web diameters of all spiders belonging to web-building guild were measured.

*S 2.3.2 Results*

In field experiments performed to investigate if spiders belonging to the web-building guild are real predators, 12 out of the 47 spiders were able to capture *O. henryi*. The difference between the web diameters of spiders that successfully predated on *O. henryi* (34.16 ± 19.81 cm, n=12) and the ones that did not (18.84 ± 13.57 cm, n=35), was significantly different (Randomisation test, P = 0.0036). Only spiders with webs bigger than about 20 cm were able to trap and capture *O. henryi*. In the second round of relative abundance sampling, performed to investigate whether web-building spiders with big webs are abundant enough to be considered as important predators of *O. henryi*, only 4 out of 106 spiders sampled had webs larger than 20 cm. Hence, spiders belonging to the web-building guild that are large enough to be capable of predating on *O. henryi* were too rare in the field and hence were not considered as major predators of *O. henryi*. Further predation risk experiments were carried out only with *P. viridans* as the main predator of *O. henryi*.

*S 2.4 Testing motivation of 24-hour starved P. viridans spiders to predate on O. henryi to determine starvation period for further experiments*

To examine whether 24-hour starvation period is enough for a large proportion of *P. viridans* (that are large enough to predate on *O. henryi*) to be motivated to predate on *O. henryi* a laboratory experiment was conducted. This allowed us to starve all spiders that were used in experiments conducted to measure probabilities of encounter and capture, for a known, short period of time.

*S 2.4.1 Methods*

59 *P. viridans* larger than the size threshold known to be capable of capturing *O. henryi* (5.12 mm) were collected in the field. These individuals were different from the ones tested in the field predation experiment to determine whether *P. viridans* are real predators of *O. henryi*. These spiders were offered *O. henryi* males inside plastic boxes in the laboratory at 1900 hours and observed every 30 minutes for a total of 3 hours to record when and whether they captured crickets. Out of the spiders that did predate on crickets, 28 were now starved for 24 hours, offered *O. henryi* males again at 1900hrs the following evening and scan sampled every 30 minutes till 2200hrs. All experiments were performed in complete darkness, similar to all other experiments.

*S 2.4.2 Results*

Out of the 59 large spiders tested in the laboratory, 40 captured crickets. Of these 40 spiders, 28 were starved for 24 hours to test for motivation to predate on crickets when starved for a short, known period of time. 24 out of 28 *P. viridans* predated on *O. henryi*. There was no significant difference between the body sizes of the spiders that captured crickets (9.51 ± 1.37 mm, n=24) and spiders that did not (9.08 ± 1.27 mm, n=4), when analysed using randomisation tests (*P* = 0.57). Hence, for all further experiments, *P. viridans* larger than 5.12 mm and starved for 48 hours, were used.

**S 3 Motivation experiment with wild-caught females**

This phonotaxis experiment was performed to determine with what proportion can wild-caught female crickets be segregated as phonotactic and non-phonotactic, as a proxy for motivation to communicate. Hence, phonotaxis was employed as a proxy for motivation to mate. Females sampled during the probability of co-occurrence study were used.

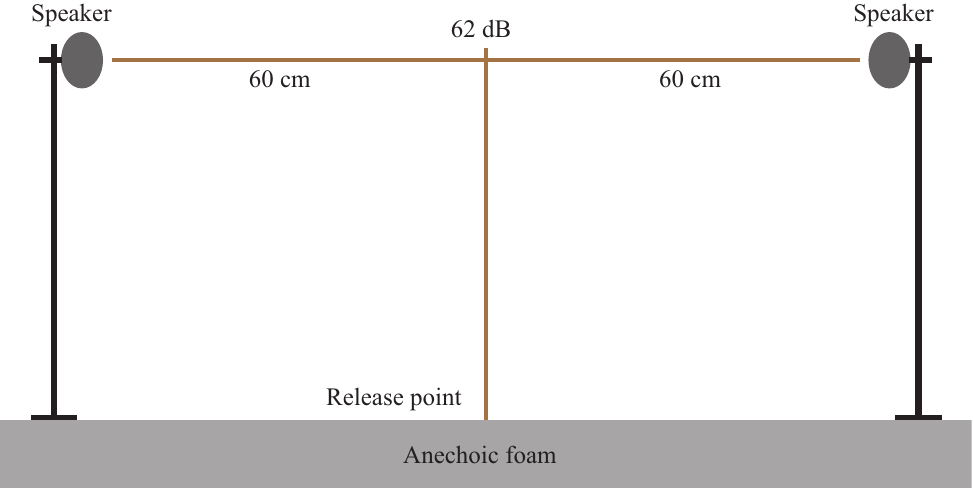

***Figure. S1.*** *Schematic representation of the phonotaxis setup used to test motivation in wild-caught O. henryi females.*

*S 3.1 Methods*

20 out of the 43 females sampled for measuring the probability of co-occurrence in the field were experimented with. The phonotaxis setup was similar to the setup described in Mhatre *et al.* (2011). It included a vertical T-shaped setup made up of stripped *H. suaveolens* branches, with each arm measuring 60 cm, placed in the centre of a room that was lined with anechoic foam (Monarch Tapes and Foams Ltd., Bangalore, India). Two speakers (X-mini Capsule Speaker V1.1, Xmi Pte Ltd, Singapore) were fixed on stands on the opposite side of the horizontal stick facing each other, without touching the horizontal stick (Fig. A1). At the start of each trial, an *O. henryi* male call was played back from one of the two speakers chosen at random such that the call was 61 dB SPL (r.m.s. re. 2 x 10^-5^ Nm^-2^) measured using a ½” microphone (Brüel and Kjær A/S, Denmark, Type 4189, 20 Hz to 20 kHz) fitted on a Sound Level Meter Type 2250 (Brüel and Kjær A/S, Denmark) at the junction of the vertical and horizontal branches, which was the decision point in the experiment. One of the 3 representative calls recorded at 22°C, 24°C and 26°C respectively, was played back depending on the temperature of the room each evening and a female was simultaneously released at the base of the T-shaped setup (calls recorded by Rittik Deb, Deb 2015). If the female reached within 10 cm of the broadcasting speaker within 300 seconds, it was considered as positive phonotaxis. These experiments were carried out between 2115 to 2145 hours.

*S 3.2 Results*

6 out of the 20 tested females performed phonotaxis. Out of the remaining 14, 6 females moved to the speaker not playing back a male call while 8 females did not move close to any of the speakers. Hence, 30% of wild-caught females were found to be motivated to perform phonotaxis towards conspecific male calls.

| Treatment | Spider in close proximity but did not attack | Spider attacked but did not capture | Spider captured cricket | Total encounters | Total number of trials |
| --- | --- | --- | --- | --- | --- |
| Calling male | 1 | 7 | 0 | 8 | 38 |
| Responding female | 4 | 10 | 2 | 16 | 45 |

Table S1. Nature of encounters between spiders and calling males and responding females along with their frequencies. For the experiment to measure the probability of encounter between crickets and spiders, an encounter was defined as any spatial proximity between the cricket and spider, within 4 cm of each other. This table details all the interactions between spiders and crickets (calling males and responding females).

**References**

Mhatre N, Bhattacharya M, Robert D, Balakrishnan R (2011) Matching sender and receiver: poikilothermy and frequency tuning in a tree cricket. Journal of Experimental Biology 214:2569–2578. doi: 10.1242/jeb.057612

Deb R (2015) Mate choice, mate sampling and baffling behaviour in the tree cricket. PhD dissertation, Centre for Ecological Sciences, Indian Institute of Science, Bangalore, India.
